# Deep immune profiling of the peripheral blood reveals disease- and sex-associated immune cell signatures in patients with systemic sclerosis

**DOI:** 10.64898/2026.05.11.724091

**Authors:** Nikhil Jiwrajka, Florin Tuluc, Nuriban Valero-Pacheco, Jennifer B. Murray, Sylvia E. Posso, Jane H. Buckner, Montserrat C. Anguera

## Abstract

**Objective:** Systemic sclerosis (SSc) predominantly affects females but exhibits greater disease severity in males, suggesting sex differences underlying SSc pathogenesis. We sought to define sex-associated alterations in the peripheral immune landscape of patients with SSc.

**Methods:** We performed high-dimensional immune profiling of PBMCs from 37 healthy donors (68% female) and 37 patients with SSc (11 limited, 26 diffuse; 68% female) using 30-color spectral flow cytometry, quantifying 56 immune cell subsets per donor. We conducted sex-stratified comparisons and correlation analysis, and used principal component analysis followed by linear discriminant analysis to derive a sex-discriminant immune cellular module.

**Results:** Diffuse cutaneous SSc (dcSSc) was associated with a distinct immune landscape characterized by increased monocyte and decreased natural killer-like and B cell frequencies, suggesting a myeloid-skewed peripheral immunophenotype. Males exhibited greater enrichment of innate immune subsets, including monocyte and dendritic cell subsets, while females exhibited greater enrichment of adaptive immune subsets. Among T cells, dcSSc was associated with coordinated remodeling across CD4^+^ and CD8^+^ subsets, including expansion of stem cell memory T cells (Tscm), and increased regulatory T cells, Th17 skewing, and decreased effector-memory CD8^+^ subsets. Females exhibited greater proportions of naïve- and Tscm, and males exhibited higher proportions of effector-memory subsets. Integrating these data, we identified a sex-discriminant immune module comprised of 20 cell types that distinguishes males and females with dcSSc.

**Conclusions:** SSc is associated with sex-specific differences in the peripheral immune landscape. A sex-associated immune program, further amplified in disease, may contribute to the paradox of female-biased susceptibility and male-biased severity in SSc.

## INTRODUCTION

Many systemic autoimmune rheumatic diseases, including systemic lupus erythematosus (SLE), systemic sclerosis (SSc), and Sjögren’s Disease, preferentially afflict females.^1^ Paradoxically, afflicted males tend to exhibit more severe disease.^2–7^ These sex differences are particularly apparent in SSc, where female-to-male prevalence ratios vary from 3:1 to 8:1.^8^ However, observational studies have consistently identified male sex as an adverse prognostic factor in SSc, with males exhibiting a greater likelihood of diffuse disease, interstitial lung disease (ILD), ILD progression, and mortality.^9,10^ These observations were further supported by a post-hoc analysis of two randomized controlled trials in SSc-ILD, which identified sex differences in baseline bronchoalveolar lavage (BAL) cytokine profiles and clinical responses to standardized immunosuppresson.^3^ Collectively, these observations suggest that sex differences in SSc susceptibility and severity may partly reflect sex differences in immune composition and/or function, though this has not been systematically investigated.

Several immune profiling studies of human peripheral blood have identified sex differences in immune cell composition and function in healthy individuals.^11–16^ Conventional flow cytometry and bulk transcriptomic approaches have identified a more prominent adaptive immune signature in females, with increased proportions of B and CD4^+^ T cells, and a more prominent innate immune signature in males, with increased proportions of monocyte and natural killer cell subsets. However, conventional cytometry is relatively limited in its ability to resolve immune heterogeneity due to constrains on number of markers that can be measured simultaneously. More recent single-cell RNA sequencing studies have enabled a high-resolution, transcriptome-wide view of sex differences across the lifespan, enabling deeper characterization of sex differences in the functional programs of immune subsets.^17–19^ While these studies have focused on healthy individuals, none, to our knowledge, have investigated whether similar or additional sex differences exist in SSc. Moreover, despite its high resolution, single-cell transcriptomics can be challenging to apply to large cohorts due to both cost and limitations on the number of cells profiled per donor, potentially reducing detection of rare immune subsets relevant to disease pathogenesis.

Spectral flow cytometry builds upon conventional flow cytometry by enabling more markers per panel with improved marker resolution. Moreover, it maintains >10-20-fold higher experimental throughput than mass cytometry and is compatible with cell sorting, enabling downstream functional and other -omics studies on purified immune cells. These capabilities render spectral cytometry well-suited to studies of immune composition and function in large patient cohorts, particularly when sample quality is limited, as a single sample can be processed for high-dimensional immunophenotyping, cell sorting, and downstream studies on purified cells. We therefore developed and applied a 30-color spectral flow cytometry panel to deeply profile peripheral blood mononuclear cell subsets from patients with SSc.

Using this platform, we performed what is, to our knowledge, the first comprehensive high-dimensional analysis of sex differences in circulating immune cells from patients with SSc. We hypothesized that, as in health, SSc would be associated with female-biased adaptive and male-biased innate immune cell profiles, and that disease-specific immune remodeling would introduce additional sex-dependent differences within individual immune subsets. We observe both disease- and sex-associated immune remodeling across innate, T cell, and B cell compartments, and identify a sex-discriminant immune cellular module that distinguishes males and females with diffuse SSc based on the relative abundance of specific immune subsets. Our findings suggest that sex-biased immune organization may contribute to the paradox of female-biased disease susceptibility and male-biased disease severity in SSc.

## METHODS

### Patient cohort and sample collection

Patients with SSc were sequentially enrolled from rheumatology clinics at Virginia Mason Medical Center into the Benaroya Research Institute Scleroderma Biorepository under an institutional review board-approved protocol. PBMCs were isolated from peripheral blood by Ficoll density gradient centrifugation and cryopreserved. Clinical data were recorded at enrollment. All patients fulfilled the 2013 ACR/EULAR classification criteria for SSc (**Supplementary Table S1)**.^20^ Patients with active malignancy, other concomitant autoimmune disease, or prior autologous stem cell or solid organ transplantation were excluded. Early disease was defined as phlebotomy within 3 years of SSc diagnosis. Healthy donors without autoimmune disease were matched to patients by sex, age, race, and duration of PBMC cryopreservation. Cryopreserved PBMCs were transferred as single vials to the University of Pennsylvania for analysis. To control for experimental batch-processing effects, PBMCs from two independent healthy donors were cryopreserved into multiple aliquots and included as technical “bridge” samples in all experimental batches.

### Spectral flow cytometry panel design, data acquisition, and analysis

A 30-color spectral flow cytometry panel was iteratively optimized to profile innate and adaptive immune populations while maximizing population resolution, minimizing artifactual spreading, and mitigating batch effects. Details regarding panel design, validation, staining and data acquisition are provided in **Supplemental Figures S1-S7**, **Supplementary Tables S2-S3**, and the **Supplementary Methods.** Fully stained samples were simultaneously acquired on a 5-Laser Cytek Aurora Analyzer and an Aurora CS. Cytometry data were analyzed using OMIQ. Following spectral unmixing, 30×30 bivariate marker plots were manually reviewed for each experimental batch to ensure unmixing accuracy. Unless specified, analyses in this study used Aurora Analyzer data. Events were manually gated to identify single, live, CD45⁺ cells for each batch. Aurora Analyzer data were then normalized using CytoNorm^21^, after which a single, fixed gating strategy was uniformly applied across all samples. Aurora CS data were used for analyzer-sorter comparisons and gated batch-specifically without normalization. Whereas downsampled data were used to facilitate UMAP-based data visualization, cell proportions were quantified using the full dataset.

### Multivariable analysis and derivation of a sex-discriminant immune module

To identify immune features associated with biological sex in diffuse SSc, cell proportion data from the 56 defined immune subsets were standardized across all dcSSc patients, and principal component analysis (PCA) was used for dimensionality reduction. Linear discriminant analysis (LDA) was applied to PCA-transformed data to identify a sex-discriminant axis. The number of principal components included in the LDA model, *k*, was selected using leave-one-out cross-validation: for each candidate value of *k*, one donor was held out, PCA was recomputed using the remaining donors, the held-out donor was projected into that PCA space, and LDA was used to classify the held-out donor by sex. Classification accuracy was calculated as the proportion of donors correctly classified across all leave-one-out iterations. The first 6 of 25 principal components were selected for the final PCA-LDA model, as this was the fewest number of components that achieved maximal classification accuracy while maintaining stable performance across adjacent *k* values.

To quantify individual immune cellular contributions to the sex-discriminant axis, LDA coefficients were back-projected into the original cell feature space using PCA loadings. Feature robustness was assessed by bootstrap resampling: in each iteration, donors were sampled with replacement, immune features were winsorized at the 5^th^ and 95^th^ percentiles to reduce the influence of outliers, and the PCA-LDA pipeline was repeated using the first 6 principal components. Feature stability was quantified based on the frequency of inclusion among top-ranked contributors to the sex-discriminant axis and consistency of effect direction across iterations. Finally, a sex-discriminant immune module score was derived from the 10 most positively weighted and 10 most negatively weighted stable cellular features by z-scaling these features across donors and computing a weighted sum using the corresponding LDA-derived feature weights.

### Statistical analyses

Statistical analyses were performed in R. Group comparisons were performed using Kruskal-Wallis tests followed by pairwise Wilcoxon rank-sum tests when appropriate. *P*-values were adjusted for multiple comparisons using the Benjamini-Hochberg method, with an adjusted *p*-value <0.05 considered statistically significant. Associations between variables were assessed using Spearman’s rank correlation coefficient with two-sided, unadjusted *P* values.

## RESULTS

### Development and Optimization of a High-Parameter Spectral Flow Cytometry Panel For Robust Immune Profiling

We developed and optimized a 30-color spectral flow cytometry panel to enable high-resolution immunophenotyping of innate and adaptive immune populations in SSc (**Supplementary Figures S1-S4**). To profile a large cohort while preserving compatibility with downstream cell sorting, we distributed cryopreserved PBMCs from 74 subjects across nine experimental batches, with each batch including aliquots from two “bridge” healthy donor samples (**Figure 1A**). Bridge samples demonstrated highly consistent immune cell distributions across batches, indicating that technical variation introduced by batch processing was substantially smaller than biological variation across donors (**Supplementary Figure S5**).

**Figure 1.**
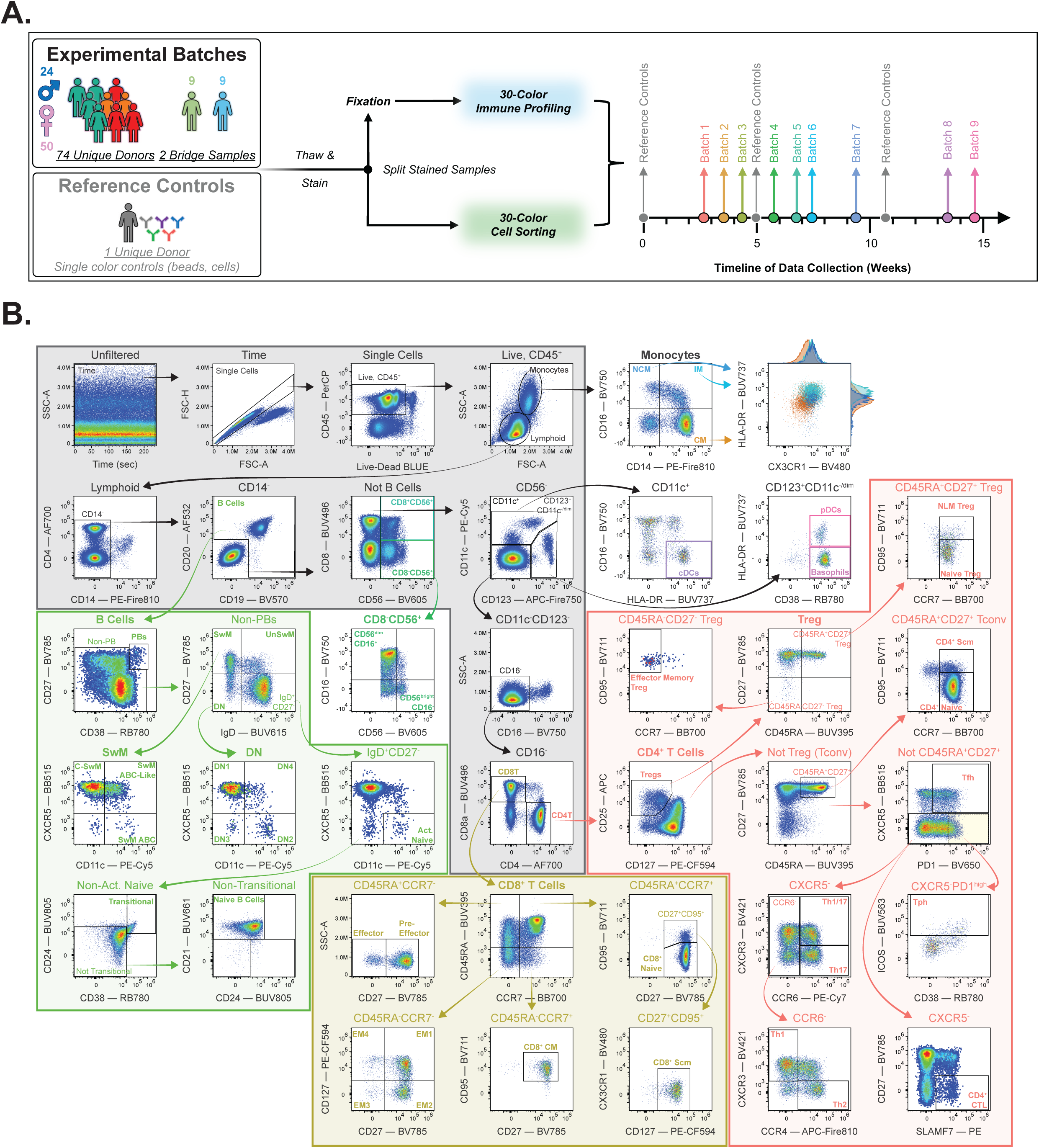
A 30-color spectral flow cytometry panel enables robust identification of diverse peripheral blood immune subsets. **A.** Overview of the experimental design. Samples from each experimental batch were processed simultaneously for high-dimensional immune profiling over a 3-month period. **B.** Bivariate gating strategy used to identify immune cell subsets of interest, shown using a representative donor sample. Cellular subset identification is as follows: monocytes (including bivariate plots to depict the relative expression of HLA-DR and CX3CR1 across monocyte subsets); B cells; CD56^+^ cells, which include natural killer (NK) and NKT cells; dendritic cells (including conventional and plasmacytoid dendritic cells); basophils; and T cells (including both CD4^+^ and CD8^+^ T cell subsets). PBs = plasmablasts; SwM = switched memory B cells; UnSwM = unswitched memory B cells; DN = double-negative B cells; Act. Naïve = activated naïve; cDCs = conventional dendritic cells; pDCs = plasmacytoid dendritic cells; Scm = stem cell memory; CM = central memory; EM1-EM4 = effector memory 1-4; Tfh = T follicular helper; Tph = T peripheral helper; CTL = cytotoxic T lymphocyte; Treg = regulatory T cell.

Following batch normalization, we applied a unified manual gating across all samples to define major immune cell populations and subsets (**Figure 1B**).^22^ We identified T cells using a negative-selection gating strategy to preserve compatibility with cell sorting for downstream activation assays. Validation of this approach using an independent dataset highlights its specificity and sensitivity for T cells: our approach captures most T cells, with >99% of the identified CD4^+^ and CD8^+^ populations expressing CD3, and recovers ≥99% of CD4^+^ T cells and ≥97% of CD8^+^ T cells identified using CD3-based gating (**Supplementary Figure S6**). Comparable population distributions were observed between analyzer- and sorter-acquired samples, supporting the utility of our panel for both high-dimensional immunophenotyping and spectral cell sorting (**Supplementary Figure S7**).

### Diffuse SSc is Associated with a Sex-Biased Peripheral Immune Landscape

We applied our panel to investigate the peripheral immune landscape of patients with SSc. Our cohort was comprised of 37 healthy donors and 37 patients with SSc, including 11 with limited cutaneous SSc (lcSSc) and 26 with dcSSc (**Supplementary Table S1**). Healthy donors and SSc patients were matched by age, sex, and race. Most SSc patients were early in their disease course, and serologic features were consistent with those expected based on SSc subtype and clinical features (**Supplementary Figure S8**). Although lcSSc patients were all female, the dcSSc subgroup was sex-balanced (14 females, 12 males), with similar exposure to immunosuppression between females and males.

We profiled 37 million live CD45^+^ cells across 74 donors (∼500,000 cells per donor). To visualize differences in immune composition, we generated UMAP projections of live, CD45^+^ cells and annotated major immune cell populations (**Figure 2A**). Comparisons of immune cell proportions revealed that patients with dcSSc exhibit a distinct peripheral immune profile relative to both healthy donors and lcSSc patients. This is characterized by increased proportions of multiple monocyte subsets, including classical (CD14^+^CD16^-^) and nonclassical (CD14^-^CD16^+^) monocytes, and by reduced CX3CR1-expressing monocytes (**Figure 2B**, **Supplementary Figure S9A**). SSc patients also exhibited lower proportions of CD8^-^ and CD8^+^ CD56^+^ lymphocyte populations, which include natural killer (NK) and NKT cells, as well as reduced total B cell frequencies. These cellular alterations were not associated with specific SSc autoantibodies, interstitial lung disease, or pulmonary hypertension (**Supplementary Figure S9B**). These findings indicate a shift toward a myeloid-skewed immune landscape in patients with diffuse disease.

**Figure 2.**
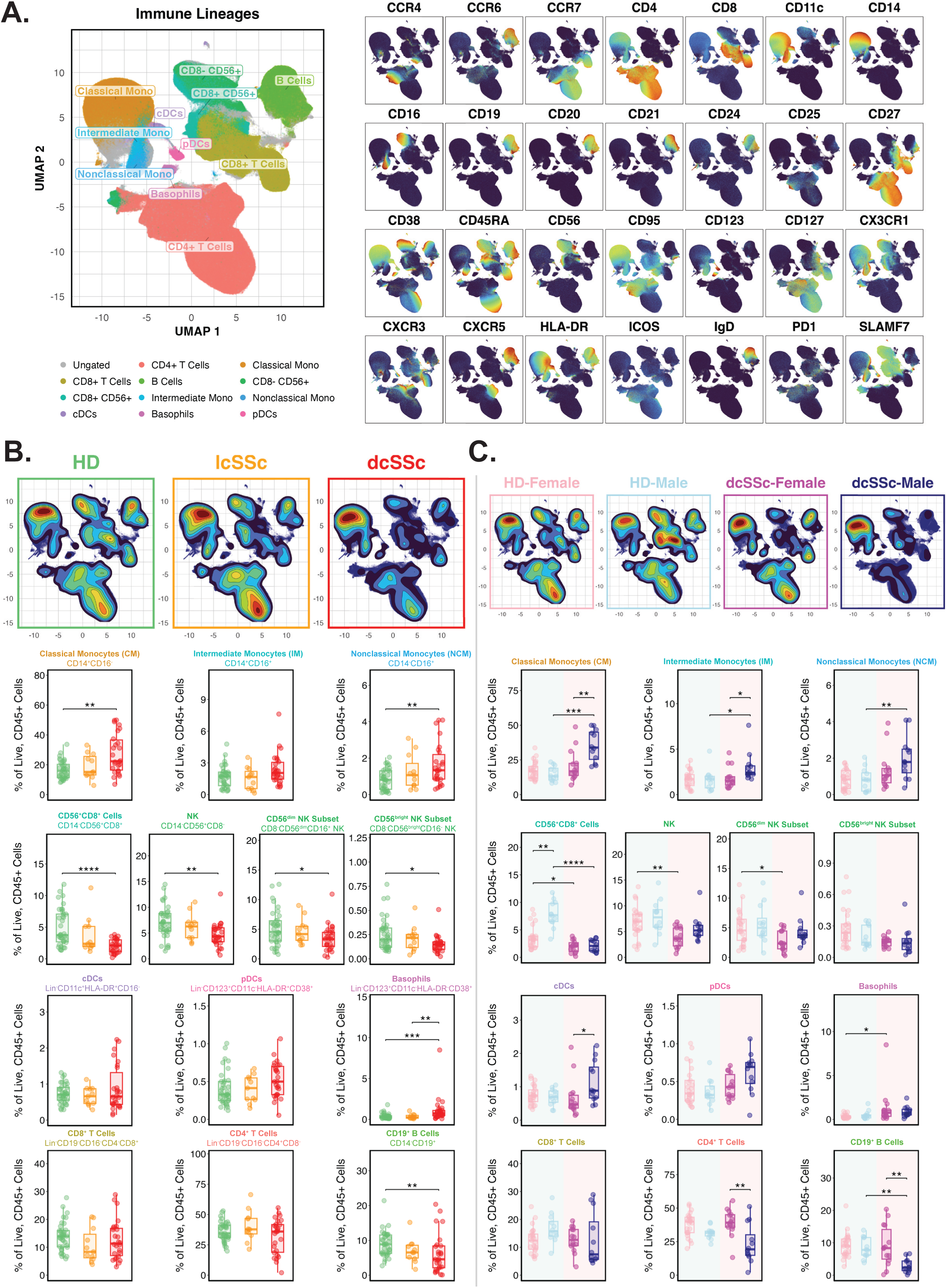
Diffuse SSc is associated with sex-biased immune profiles spanning innate and adaptive immunity. **A.** UMAP projection of live, CD45^+^ cells from healthy donors (n=37), patients with limited cutaneous SSc (lcSSc; n=11), and patients with diffuse cutaneous SSc (dcSSc; n=26), annotated according to the gating strategy depicted in Figure 1B, and constructed using all markers except CD45-PerCP. A total of 1.5 million cells is shown, downsampled from a total of ∼37 million live, CD45^+^ cells, with equal event sampling per donor within each group (∼13,000-45,000 cells per donor depending on the group). Batch-normalized expression of the remaining 28 panel markers is depicted, overlaid on the UMAP embedding. **B.** Contour density plots of group-specific UMAP projections and quantified proportions of distinct immune lineages across disease states, shown as boxplots. **C.** Sex-stratified contour density plots of UMAP projections for healthy donors and patients with dcSSc and quantified proportions of distinct immune lineages stratified by sex, shown as boxplots. *, **, ***, or **** = adjusted *p* < 0.05, 0.01, 0.001, or 0.0001, respectively, via Kruskal-Wallis test with Benjamini-Hochberg adjustment for multiple comparisons.

We subsequently performed sex-stratified analyses of these immune cell populations. Many of the innate immune cell expansions observed in dcSSc were predominantly driven by affected males (**Figure 2C**). In addition to increased monocyte subsets, males with dcSSc exhibited increased conventional dendritic cells, with a similar trend for plasmacytoid dendritic cells. Conversely, females exhibited relatively higher proportions of adaptive immune populations, including CD4^+^ T cells and B cells (**Figure 2C**), indicating that the broad immune shifts identified in patients with diffuse disease do not occur uniformly across the sexes. Notable exceptions included CD56^+^CD8^+^ cells, which were enriched in healthy males relative to females but were uniformly reduced in both sexes in dcSSc (**Figure 2C**). In contrast, CD56^+^CD8^-^ cells showed minimal sex differences in health but were also decreased in dcSSc regardless of sex. Collectively, these data suggest that the peripheral immune landscape of dcSSc is strongly shaped by biological sex, with males exhibiting an innate-skewed profile and females exhibiting relative enrichment of adaptive immune populations.

### CD8^+^ T Cells in SSc Exhibit Female-Biased Enrichment of Stem-Like Memory Cells and Sex-Skewed Effector Memory Differentiation

To further resolve alterations within adaptive immunity, we first examined the CD8^+^ T cell compartment across disease states. UMAP visualization and quantitative analysis revealed that both lcSSc and dcSSc were characterized by increased proportions of CD8^+^ stem cell memory (Tscm) T cells, a rare memory subset expanded in other systemic autoimmune diseases including SLE and RA (**Figure 3A-B)**.^23–25^ Both lcSSc and dsSSc also exhibited a reduction in effector-memory subsets (**Figure 3A-B**), potentially reflecting accumulation of these cells in lesional skin.^26^ Although CD8^+^ T cell subsets were not strongly associated with specific clinical features or autoantibodies, several subsets exhibited expected age-related changes^18,19,27^, including declining total and naïve CD8^+^ T cells with age, and a relative increase in more differentiated memory populations (**Figure 3C**, **Supplementary Figure S10**).

**Figure 3.**
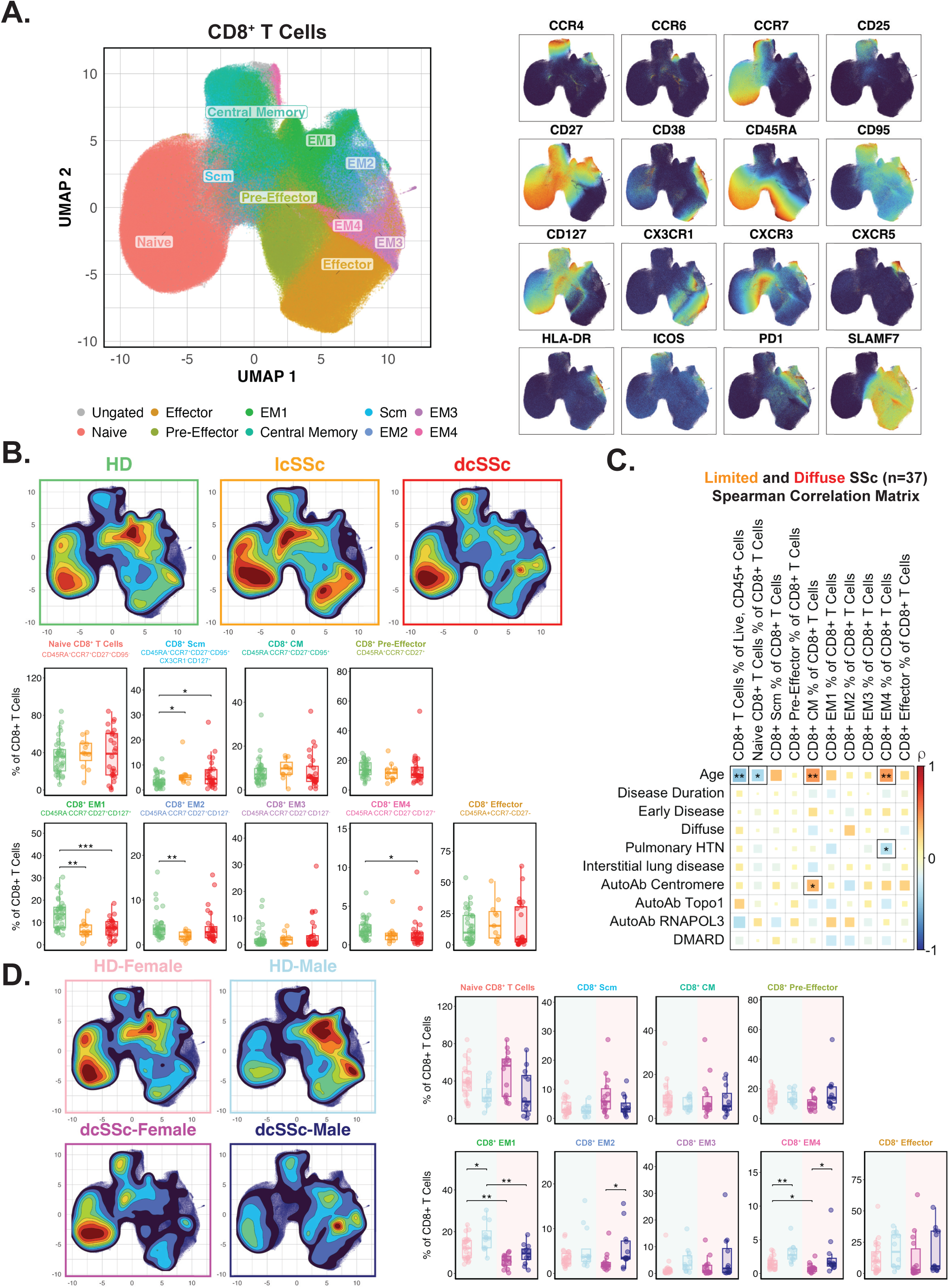
SSc is associated with reduced effector-memory CD8^+^ T cells and sex-biased CD8^+^ T cell differentiation. **A.** UMAP projection of CD8^+^ T cells from healthy donors, patients with lcSSc, and patients with dcSSc, annotated according to the gating strategy depicted in Figure 1B, and constructed using 16 markers to identify CD8^+^ T cell subsets. A total of 955,000 cells is depicted, downsampled from a total of 4.8 million CD8^+^ T cells, with equal event sampling per donor within each group (∼10,000-15,000 cells per donor depending on the group). Batch-normalized expression of the 16 markers used to construct the UMAP is depicted, overlaid on the UMAP embedding. **B.** Contour density plots of group-specific UMAP projections and quantified proportions of distinct subsets across disease states, shown as boxplots. **C.** Spearman (rho) correlation matrix of clinical features and CD8^+^ T cell subsets among all SSc patients (n=37). Statistically significant correlations are indicated with a solid black frame and at least one asterisk. *, ** = unadjusted *p* < 0.05 or 0.01, respectively, based on Spearman’s rank correlation coefficient. **D.** Sex-stratified contour density plots of UMAP projections for healthy donors and patients with dcSSc and quantified proportions of distinct subsets stratified by sex, shown as boxplots. For boxplots in panels B and D, *, **, or *** = adjusted *p* < 0.05, 0.01, or 0.001, respectively, via Kruskal-Wallis test with Benjamini-Hochberg adjustment for multiple comparisons.

Sex-stratified analyses revealed sex-associated divergence across the CD8^+^ differentiation landscape. In both health and disease, females tended to exhibit greater proportions of naïve and Tscm CD8^+^ T cells, while males exhibited relatively higher proportions of effector-memory subsets (**Figure 3D**). Notably, although effector-memory populations were reduced overall in dcSSc, they remained relatively enriched in affected males compared to females. Given the profibrotic role of effector-memory CD8^+^ T cells in SSc skin^28^, these findings suggest that male-biased enrichment of these subsets may contribute to sex-biased inflammation in lesional tissues. Together, these data suggest that CD8^+^ T cell differentiation is shaped by both disease status and sex, with males exhibiting a more differentiated effector-memory profile and females retaining a more naïve and stem-like phenotype.

### CD4^+^ T Cells in Diffuse Disease Exhibit Sex-Biased Variation in Regulatory, Naïve, and Stem Cell Memory T Cells

CD4^+^ T cell subsets have long been implicated in SSc pathogenesis, particularly through their roles in proinflammatory and profibrotic cytokine production and B cell help.^29^ We therefore examined the CD4^+^ T cell compartment by disease state. Patients with dcSSc exhibited increased proportions of regulatory T cells (Tregs), including expansion of a naïve-like memory Treg subset^30^ with increased suppressive capability and greater clonality relative to naïve Tregs (**Figure 4A-B**). As observed with CD8^+^ T cells, patents with dcSSc also exhibit increased proportions of conventional (non-Treg) CD4^+^ Tscm, along with a modest increase in the proportion of T peripheral helper (Tph; CXCR5^-^PD1^high^ICOS^+^) cells (**Figure 4B**). In contrast, overall proportions of naïve CD4^+^ T cells, T follicular helper (Tfh), and cytotoxic CD4^+^ T cells were not significantly altered. Analysis of T helper polarization revealed reduced Th1-polarized and expanded Th17-polarized CD4^+^ T cells in dcSSc, with no significant differences in Th2-polarized cells (**Figure 4B**). Associations between CD4⁺ T cell subsets and clinical features were generally modest (**Figure 4C**). Th17-polarized cells were positively associated with pulmonary hypertension, as previously reported.^31^ Cross-lineage correlation with CD8^+^ T cell subsets revealed synchronous changes in naïve and non-naïve subsets, suggesting coordinated remodeling across T cell compartments (**Supplementary Figure S11**).

**Figure 4.**
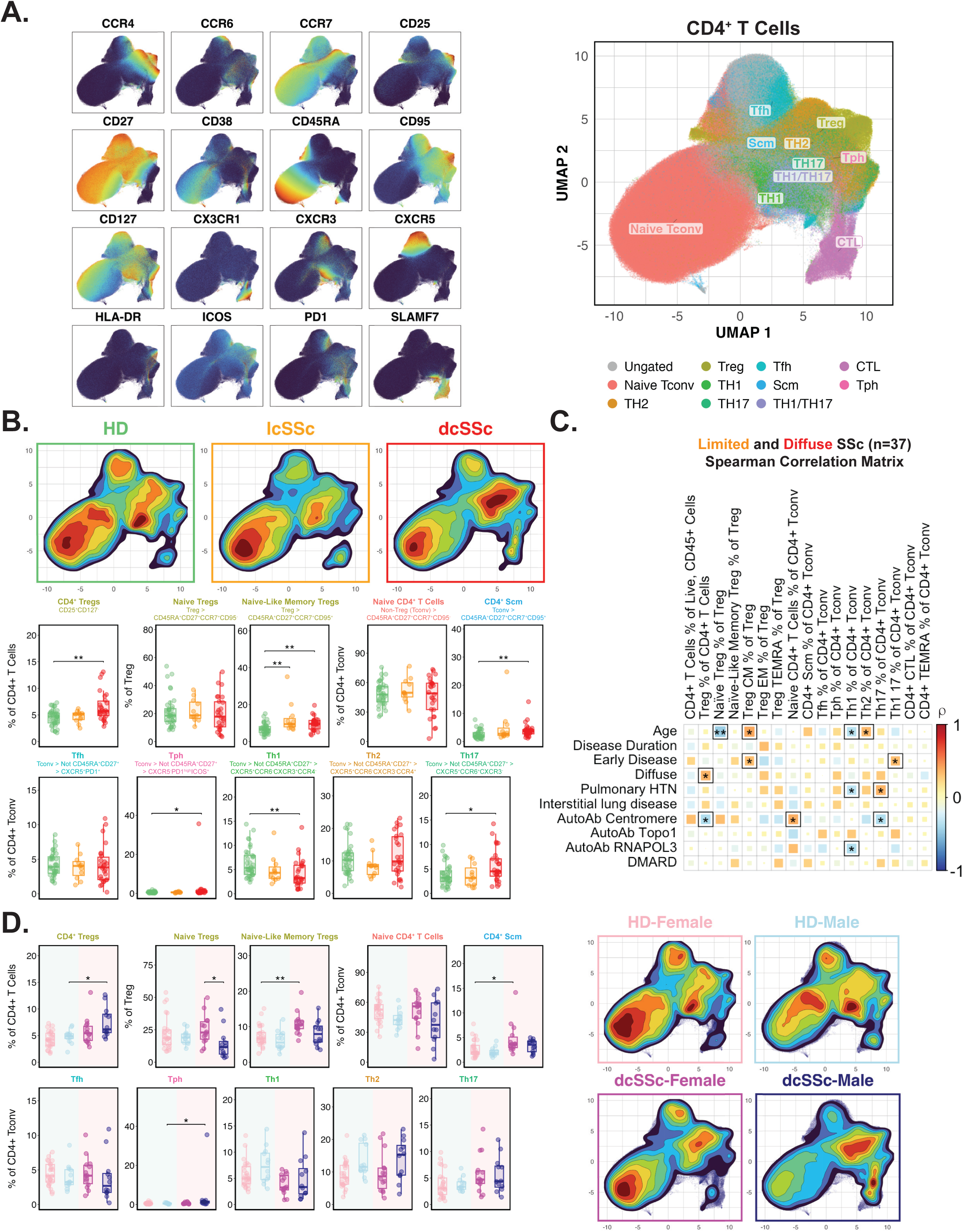
CD4^+^ T cell remodeling in diffuse SSc includes regulatory T cell expansion, Th17-skewing, and sex-biased enrichment of naïve and stem-like memory subsets. **A.** UMAP projection of CD4^+^ T cells from healthy donors, patients with lcSSc, and patients with dcSSc, annotated according to the gating strategy depicted in Figure 1B, and constructed using 16 markers to identify CD4^+^ T cell subsets. A total of ∼908,000 cells is shown, downsampled from a total of 13 million CD4^+^ T cells, with equal event sampling per donor within each group (∼9,000-30,000 cells per donor depending on the group). Batch-normalized expression of the 16 markers used to construct the UMAP is depicted, overlaid on the UMAP embedding. **B.** Contour density plots of group-specific UMAP projections and quantified proportions of distinct immune subsets across disease states, shown as boxplots. **C.** Spearman (rho) correlation matrix of clinical features and CD4^+^ T cell subsets among all SSc patients (n=37). Statistically significant correlations are indicated with a solid black frame and at least one asterisk. *, ** = unadjusted *p* < 0.05 or 0.01, respectively, based on Spearman’s rank correlation coefficient. **D.** Sex-stratified contour density plots of UMAP projections for healthy donors and patients with dcSSc and quantified proportions of distinct immune subsets stratified by sex, shown as boxplots. For boxplots in panels B and D, *, ** = adjusted *p* < 0.05, 0.01, respectively, via Kruskal-Wallis test with Benjamini-Hochberg adjustment for multiple comparisons.

Sex-stratified analyses revealed sex-associated heterogeneity within the CD4⁺ compartment. Whereas overall Treg and Tph expansion in dcSSc was primarily observed in affected males, females tended to exhibit greater proportions of naïve and naïve-like memory Tregs (**Figure 4D**). Consistent with our findings in CD8^+^ T cells, females also tended to exhibit greater proportions of naïve CD4^+^ and CD4^+^ Tscm cells (**Figure 4D**). In contrast, disease-associated shifts in T helper polarization were largely conserved across sexes (**Figure 4D).** Overall, CD4^+^ T cell remodeling in dcSSc parallels patterns observed in CD8^+^ T cells, with afflicted females exhibiting coordinated enrichment of naïve and stem-cell-memory-like subsets.

### Memory B Cell Remodeling in dcSSc Reflects a Largely Sex-Independent Shift Toward CD11c^+^ Subsets

Given the encouraging results of targeted B cell depletion in SSc and the association between disease-specific autoantibodies and SSc phenotypes, we examined the peripheral B cell compartment.^32,33^ Patients with dcSSc exhibited increased proportions of circulating plasmablasts and modest shifts in naïve and transitional subsets (**Figure 5A-B**). In contrast, we observed marked shifts within the class-switched (SwM; IgD^-^CD27^+^) and double-negative (DN; IgD^-^CD27^-^) memory B cell compartments in patients with dcSSc. SwM B cells exhibited a redistribution away from canonical SwM (CXCR5^+^CD11c^-^) towards CD11c-expressing populations, including age-associated B cells (“ABC”; CXCR5^-^CD11c^+^) and ABC-like (CXCR5^+^CD11c^+^) subsets (**Figure 5B**). Similarly, within DN B cells, we observed a contraction of DN1 (CXCR5^+^CD11c^-^) cells and expansion of both DN2 (CXCR5^-^CD11c^+^) and DN3 (CXCR5^-^CD11c^-^) subsets. Among SSc patients, CXCR5^-^CD11c^+/-^ subsets, including activated naive, SwM ABC, DN2, and DN3 cells, were strongly correlated with one another (**Supplementary Figure S12A**). Given the role of CD4^+^ T cells in shaping humoral responses, we also examined relationships between the T and B cell compartments. We observed strong correlations between circulating Tph, but not Tfh, cells and CXCR5^-^CD11c^+/-^ memory B cell subsets, including SwM ABC, DN2, and DN3 B cells, consistent with prior observations in SLE^34^, suggesting that Tph functions may shape coordinated B cell remodeling in SSc (**Figure 5C**, **Supplementary Figure S12B-C**).

**Figure 5.**
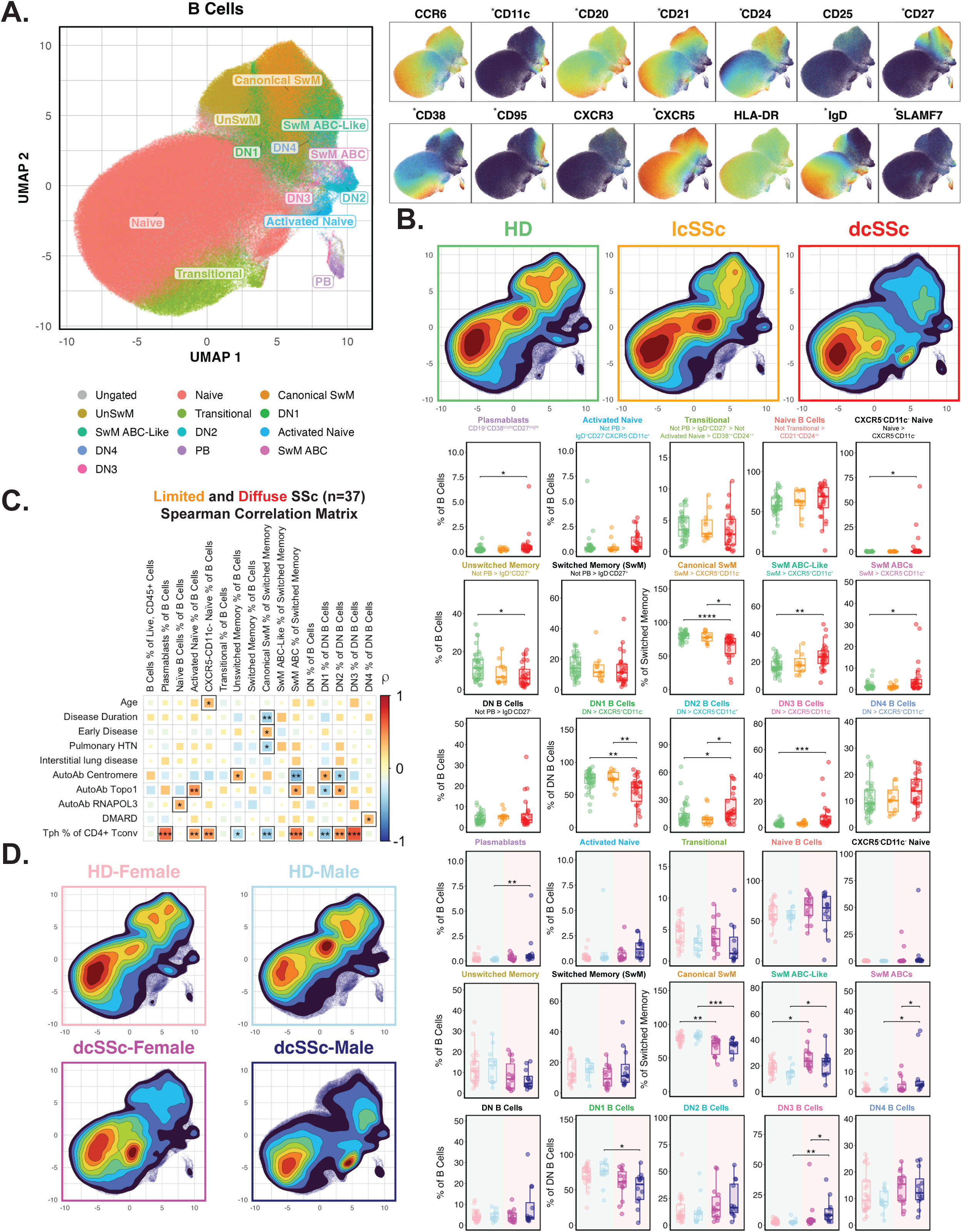
Memory B cells from patients with diffuse SSc are enriched for CD11c-expressing subsets. **A.** UMAP projection of CD19^+^ B cells from healthy donors, patients with lcSSc, and patients with dcSSc, annotated according to the gating strategy depicted in Figure 1B, and constructed using 10 markers (asterisked) to identify CD19^+^ B cell subsets. A total of ∼475,000 cells is shown, downsampled from a total of 2.8 million CD19^+^ B cells, with equal event sampling per donor within each group (∼4,000-15,000 B cells per donor depending on the group). Batch-normalized expression of 14 markers is depicted, overlaid on the UMAP embedding. **B.** Contour density plots of group-specific UMAP projections and quantified proportions of distinct immune subsets across disease states, shown as boxplots. **C.** Spearman (rho) correlation matrix of clinical features and CD19^+^ B cell subsets among all SSc patients (n=37). Statistically significant correlations are indicated with a solid black frame and at least one asterisk. *, **, *** = unadjusted *p* < 0.05, 0.01, or 0.001, respectively, based on Spearman’s rank correlation coefficient. **D.** Sex-stratified contour density plots of UMAP projections for healthy donors and patients with dcSSc and quantified proportions of distinct immune subsets stratified by sex, shown as boxplots. For boxplots in panels B and D, *, **, *** = adjusted *p* < 0.05, 0.01, or 0.001, respectively, via Kruskal-Wallis test with Benjamini-Hochberg adjustment for multiple comparisons.

Associations between B cell subset and clinical features suggest distinct immune pathways underlying disease heterogeneity. CD11c^+^ memory B cell subsets, including DN2 and SwM ABCs, were positively correlated with anti-topoisomerase I autoantibodies, and negatively correlated with anti-centromere autoantibodies, whereas unswitched memory B cells showed the opposite pattern (**Figure 5C**). These findings suggest that distinct B cell differentiation trajectories may contribute to specific autoantibody profiles in SSc.

In contrast to T cells, diffuse disease-associated changes in B cell subsets were largely conserved across the sexes **(Figure 5D**). Of the four CD11c^+^ memory subsets (DN2, DN4, SwM SwM ABC-Like, and SwM ABCs), only SwM ABCs exhibited sex-biased enrichment, with greater proportions in affected males (**Figure 5D).** Thus, SSc is associated with considerable reorganization of the memory B cell compartment, with expansions of CXCR5^-^CD11c^+^ SwM and DN subsets that are associated with T cell help, autoantibody specificity, and both germinal center and extrafollicular origins.

### A Sex-Discriminant Immune Module Distinguishes Males and Females with dcSSc

Having quantified 56 immune subsets per donor, we next asked whether these immune profiles could distinguish patients by disease status and sex. Principal component analysis (PCA) across all 74 donors showed that PC1 and PC2 partially distinguished dcSSc from lcSSc and healthy donors; however, lcSSc largely overlapped with healthy donors (**Figure 6A**). Examination of cell feature loadings indicated that the dcSSc-enriched region of PCA space was driven by innate immune populations, including monocyte subsets, as well as CD4^+^ Tscm and CD11c^+^ memory B cell subsets (**Figure 6B**), consistent with our lineage-stratified analyses.

**Figure 6.**
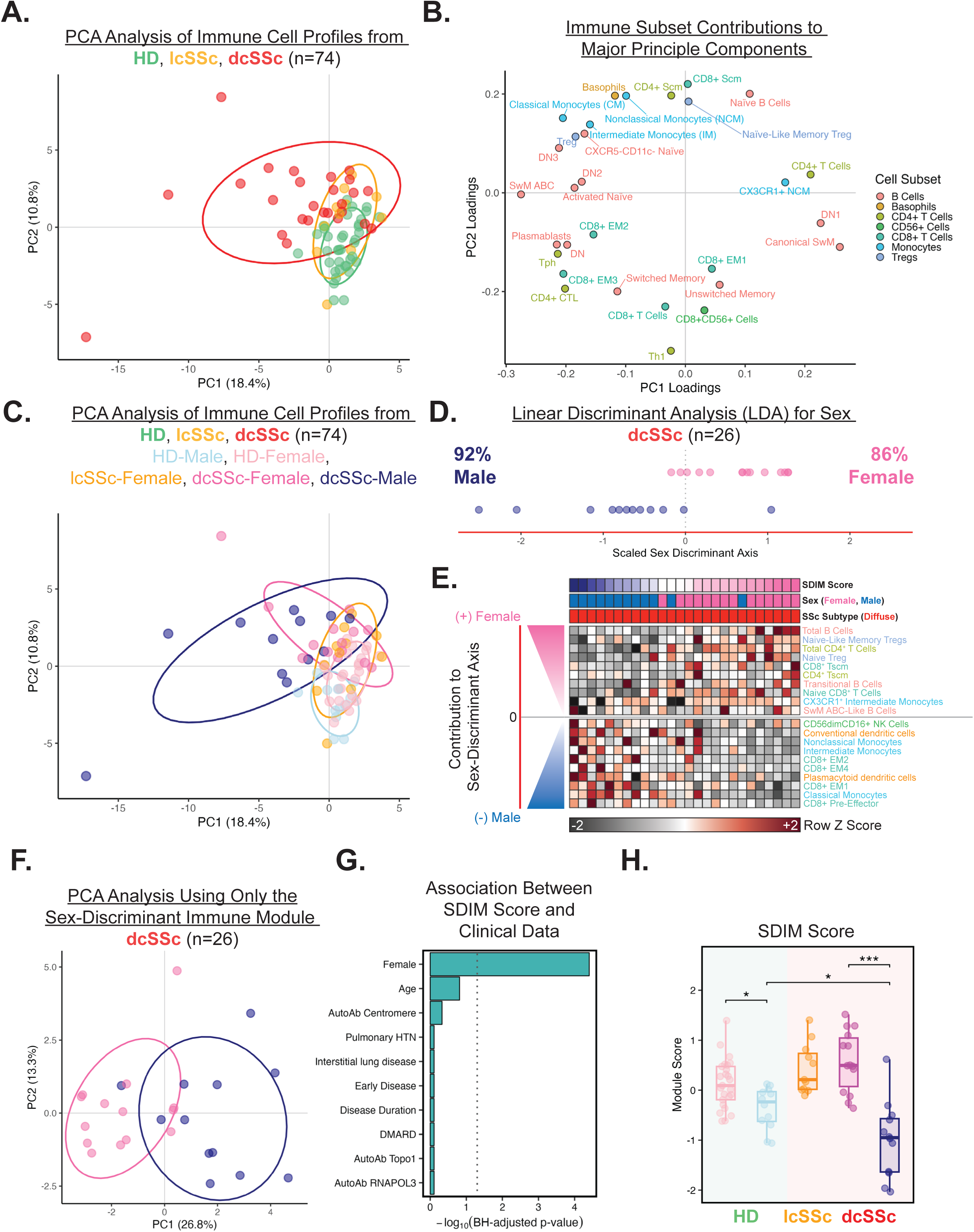
Integration of immune cell profiles identifies a sex-discriminant immune module that distinguishes males and females with diffuse SSc. **A.** Principal component analysis (PCA) of peripheral blood immune profiles from healthy donors, patients with lcSSc, and patients with dcSSc, colored by disease status. Immune profiles were comprised of the 56 enumerated cell subsets, and each point represents one donor. **B.** Top 30 cellular features contributing to principal components 1 and 2. **C.** PCA plot from panel A, colored by disease status and sex. **D.** Scaled sex-discriminant axis derived from linear discriminant analysis of immune profiles from patients with dcSSc (n=26). **E**. Heatmap of patients with dcSSc depicting the 10 strongest stable cellular contributors to each end of the sex-discriminant axis. **F.** PCA of patients with dcSSc using the 20-cell sex-discriminant immune module. **G.** Associations between the sex-discriminant immune module (SDIM) score and clinical variables among patients with dcSSc. **H**. Boxplots of SDIM scores across sex-stratified disease subgroups. *, **, ***, **** = adjusted *p* < 0.05, 0.01, 0.001, 0.0001, respectively, via Kruskall-Wallis test with Benjamini-Hochberg adjustment for multiple comparisons.

The dominant principal components did not readily distinguish females and males with dcSSc (**Figure 6C**), suggesting that sex-associated variation may not align with the major axes of overall immune variation. We therefore applied linear discriminant analysis (LDA) to PCA-transformed data from patients with dcSSc to identify a distinct sex-discriminant axis that separated dcSSc females from dcSSc males (**Figure 6D**). We subsequently generated a 20-cell sex-discriminant immune module (SDIM) comprising the 10 strongest cellular contributors to each end of the sex-discriminant axis (**Figure 6E**). The female end of this axis was driven by naïve and stem-like lymphocyte populations, whereas the male end was driven by innate and effector-memory CD8^+^ T cells. These 20 immune cell types were sufficient to distinguish males and females with dcSSc (**Figure 6E-F**). By taking a weighted sum of their contributions to the sex-discriminant axis, we derived a SDIM score, with high scores associated with female sex and low scores with male sex (**Figure 6E**). This module score was not strongly associated with other clinical variables, suggesting its specificity for biological sex among dcSSc patients **(Figure 6G**). When applied across the full cohort, the SDIM score also distinguished healthy males and females, indicating that this sex-associated immune program may be present in health but is further magnified in diffuse disease (**Figure 6H**). Collectively, these findings identify a unique immune signature that captures sex-specific variation in SSc, and suggests that differential immune organization in males and females may contribute to the sexually divergent clinical trajectories observed in patients.

## DISCUSSION

We developed and applied a robust spectral cytometry platform to deeply profile PBMCs from healthy donors and patients with SSc. We found that patients with diffuse, but not limited, SSc exhibit a distinct circulating immune profile characterized by expansion of monocyte and dendritic cell subsets, reduction of NK and effector-memory CD8^+^ T cell subsets, enrichment of stem-cell-memory and Th17-polarized T cells, and shifts within the memory B cell compartment towards CD11c^+^ subsets. Several of these changes were strongly associated with biological sex: females with dcSSc exhibited relative enrichment of lymphocyte subsets, particularly naïve and stem cell memory T cells, whereas males exhibited enrichment of myeloid subsets. Integrating these features, we identified a sex-discriminant immune cellular module that distinguished females and males both in health and disease. Our work provides a high-resolution view of peripheral immune organization in SSc and the first detailed investigation of sex differences in immune composition in SSc, enabling new insights into immunologic factors that may contribute to sex biases in SSc susceptibility and severity.

Our findings are broadly consistent with prior studies in SSc describing expansion of circulating monocyte subsets^35,36^, altered NK cell^37^ and pDC^35,38^ abundance, increased Tregs and Th17-polarized CD4⁺ T cells^31,39–42^, and changes in CD8⁺ effector-memory subsets^43^. However, peripheral blood measurements reflect a dynamic equilibrium between circulating and tissue-infiltrating immune cells; thus, differences in cell abundance may reflect altered trafficking, systemic expansion or contraction, or both. This is particularly relevant in SSc, where several populations altered in the circulation, including monocytes^44,45^, NK cells^46,47^, pDCs^38^, and effector-memory CD8^+^ T cells^26^, have also been described within affected skin or lung tissues. These observations support the biological relevance of the immune alterations identified here while underscoring the need for future paired peripheral and tissue-level analyses.

Beyond corroborating established patterns, our data highlight several less well-characterized immune populations that may contribute to SSc pathogenesis. First, we identified expansion of Tscm cells across both CD4^+^ and CD8^+^ compartments. Although Tscm have been described in RA and SLE, their potential role in SSc has not been well-defined.^48,49^ Given their capacity to serve as durable, self-renewing sources of T cell memory, Tscm may represent reservoirs of autoreactive lymphocytes capable of sustaining chronic T cell activation in the setting of persistent autoantigen exposure. Future studies to ascertain Tscm clonal diversity and whether these clones are capable of recognizing SSc autoantigens could help determine their role in SSc pathogenesis. Second, we observed a coordinated shift across the memory B cell compartment in dcSSc toward CD11c^+^ subsets (SwM ABC-Like, SwM ABC, DN2) and CD11c^-^DN3 B cells. Several subsets, including DN2 B cells, were positively correlated with anti-topoisomerase I autoantibodies. A broader population (IgD^-^CD27^-^CD21^-/lo^) encompassing both DN2 and DN3 B cells was previously found to be enriched within the circulation and the lungs of patients with SSc-ILD.^50^ While typically observed transiently in the circulation after antigen exposure, these memory B cell subsets may exhibit aberrant persistence in the setting of chronic autoantigen exposure and contribute to disease, as has been suggested in SLE^51^ and IgG4-related disease^52^. Interestingly, these subsets were strongly correlated with circulating Tph cells, which arise in chronic type 1 interferon-conditioned environments^53,54^ and promote peripheral B cell activation partly via IL-21, which induces CD11c expression in B cells.^55^ Together with prior studies identifying Tph cells in SSc skin^56^, these findings suggest that coordinated Tph-B cell interactions may contribute to the expansion or persistence of these atypical memory B cell subsets in SSc.

A central goal of our study was to determine how biological sex shapes immune composition in SSc. Our integrated analyses revealed that sex-associated variation in SSc was organized across multiple immune compartments rather than any single immune lineage, highlighting the value of a high-throughput methodology capable of profiling broadly across immune lineages while retaining sufficient depth to resolve less abundant subsets across many donors. Our sex-discriminant immune cell module reduced this high-dimensional variation to a defined set of 20 innate and adaptive immune subsets that distinguished males and females in both disease and health. The male end of this module was enriched for monocyte and dendritic cell subsets^35,38,44,45^, and effector-memory CD8^+^ T cells^26,47^—populations implicated in profibrotic pathways in SSc. In contrast, the female end was enriched for B cells, naïve T cells, and Tscm, suggesting relative enrichment of lymphocyte populations capable of broad antigen responsiveness and durable memory. These findings are consistent with established patterns of sex-associated immune variation in healthy individuals^57^, and suggest that female-biased enrichment of naïve and stem-like lymphocyte pools may provide immunological infrastructure for heightened autoreactive potential, whereas male-biased enrichment of innate and effector populations may enable more tissue-directed fibroinflammatory programs. Notably, our findings may also help reconcile prior observations in SSc-ILD. In a post-hoc analysis of a randomized trial comparing mycophenolate mofetil (MMF) with cyclophosphamide, males experienced worse outcomes than females, including slower improvement with MMF and increased mortality after adjustment for confounding factors.^3^ Whole-blood transcriptional analyses from MMF-treated patients showed that higher baseline myeloid module scores were associated with worse pulmonary outcomes, whereas higher lymphoid module scores predicted improved pulmonary function. Although the transcriptional analyses were not sex-stratified, their association with distinct clinical trajectories is consistent with the male-biased myeloid and female-biased lymphoid cellular patterns identified here.

The observed sex differences in immune organization likely arise from coordinated effects of sex hormones and genetic mechanisms. Androgens suppress thymopoiesis and B cell lymphopoiesis, whereas estrogens can promote autoreactive B cell survival by impairing central and peripheral tolerance checkpoints, providing potential mechanisms for sex-biased lymphocyte abundance.^58–64^ However, recently identified sex-specific autosomal susceptibility loci in SSc indicate that biological sex also influences disease pathogenesis at the genetic level.^65^ In parallel, variable escape from X-chromosome inactivation may increase expression of select X-linked immunomodulatory genes in female immune cells^66^, including genes such as *KDM6A* that regulate immune cell abundance, function, and pathways relevant to stem-like T cell states.^67,68^ Importantly, mechanisms driving sex differences in immune composition may also alter immune activation and effector function, underscoring the need for future studies integrating immune phenotyping with functional analyses.

Notably, many immune alterations identified here were most pronounced in dcSSc, with fewer differences observed in lcSSc. This may reflect true SSc subtype-specific differences, as several previously described alterations—including changes in circulating and/or skin-infiltrating NK cells, pDCs, Th17-polarized cells, Tph cells, and CD8^+^ effector-memory subsets—have been most prominently observed in early dcSSc. However, this interpretation is complicated by the epidemiology and heterogeneity of SSc subtypes: lcSSc is more strongly female-biased, while dcSSc is generally more severe and relatively enriched for males. Consistent with this overlap between SSc subtype and sex, SDIM scores in patients with lcSSc, all of whom were female, were similar to those observed in healthy females and females with dcSSc, and distinct from those observed in males. Thus, the limited differences observed in lcSSc may reflect subtype-specific immune biology, the strongly female-skewed composition of this subgroup, and/or immune variation driven by other features including disease severity and extracutaneous involvement. Larger cohorts with balanced sex representation across disease subgroups and detailed clinical phenotyping could help clarify whether limited and diffuse SSc are characterized by distinct immune programs or shared programs that differ in magnitude.

Our study has several limitations. The cohort did not include systematically collected measures of disease severity, limiting our ability to define relationships between specific immune subsets and clinically severe disease, particularly in sex-stratified analyses, and potentially impacting our inability to resolve distinct immune profiles in patients with lcSSc. Second, SSc patients in this cohort were receiving immunosuppressive therapy; although the frequency of immunosuppression was similar across SSc subgroups and between females and males with dcSSc, treatment exposure may have affected our comparisons between healthy donors and SSc patients. Despite these limitations, this study has important strengths, including use of a rigorously optimized spectral cytometry platform enabling high-throughput quantification of a broad range of immune subsets. To our knowledge, our study represents one of the most comprehensive immunophenotypic analyses in SSc to date and the first to systematically define sex-associated variation across multiple immune lineages. Our data provide a foundation for future studies to validate these sex-associated immune programs, define their underlying hormonal and genetic mechanisms, and determine how they differentially impact immune function in SSc, ultimately enabling insights into novel factors that may shape the clinical trajectories of patients with SSc.

## SUPPLEMENTARY TABLES

**Supplementary Table 1.** Patient characteristics. Percentages are calculated based on columns. * = Denominator for percentage calculation reflects missing data. ^†^ = DMARD includes methotrexate (n= 13), mycophenolate mofetil (n= 7), or cyclophosphamide (n=1).

|  | Healthy Donor |  | Systemic Sclerosis (SSc) |  |  |
| --- | --- | --- | --- | --- | --- |
|  |  |  | lcSSc | dcSSc |  |
| Total n (% of group) | 37 (100.0%) |  | 11 (29.7%) | 26 (70.3%) |  |
| Sex (n, %) | <u>Female</u><br>(25, 67.6%) | <u>Male</u><br>(12, 32.4%) | <u>Female</u><br>(11, 100.0%) | <u>Female</u><br>(14, 53.8%) | <u>Male</u><br>(12, 46.2%) |
| Age (mean ± S.D.) | 50.7 ± 11.5 | 53.3 ± 10.4 | 57.9 ± 11.1 | 48.1 ± 11.2 | 54.3 ± 10.5 |
| White (n, %) | 21, 84.0% | 11, 91.7% | 9, 90.0%* | 10, 76.9%* | 8, 100.0%* |
| Median Disease Duration in Years [IQR] | - | - | 0.3 [0.1 - 1.1] | 2.5 [1.7 - 8.8] | 2.2 [0.5 - 4.4] |
| Early Disease<sup>†</sup> (n, %) | - | - | 9 (81.8%) | 9 (64.3%) | 7 (58.3%) |
| Interstitial Lung Disease (n, %) | - | - | 5 (45.5%) | 7 (50.0%) | 6 (50.0%) |
| Pulmonary Hypertension (n, %) | - | - | 1 (9.1%) | 5 (35.7%) | 2 (16.7%) |
| Anti-Centromere Autoantibodies (n, %) | - | - | 5* (55.6%) | 2* (15.4%) | 0* (0.0%) |
| Anti-Topoisomerase I Autoantibodies (n, %) | - | - | 5* (55.6%) | 11* (84.6%) | 7* (70.0%) |
| Anti-RNAPOL3 Autoantibodies (n, %) | - | - | 0* (0.0%) | 0* (0.0%) | 2* (20.0%) |
| DMARD<sup>†</sup> (n, %) | - | - | 5 (45.5%) | 8 (57.1%) | 7 (58.3%) |

**Supplementary Table S2.** List of antibody-fluorophore conjugates used in this panel.

**Supplementary Table S3.** List of additional cytometry reagents used in this panel.

## SUPPLEMENTARY FIGURES

**Supplementary Figure 1. Stepwise approach to fluorochrome and antibody-fluorochrome conjugate selection for immunophenotyping human peripheral blood mononuclear cells.**

This panel was created in three broad steps, starting with an 18-marker panel to profile CD4^+^ and CD8^+^ T cell subsets. Additional lineage-defining markers, followed by B cell subset markers, were sequentially added to produce the final 30-marker panel. Fluorochrome selection was guided in part by manual review of fluorochrome spectral overlap, analyses of pairwise similarity indices, and overall panel Complexity Indices. **A**. Heatmap of similarity indices of an initial 18-marker panel designed to profile CD4^+^ and CD8^+^ T cells, as calculated in Cytek® Full Spectrum Viewer. **B.** Heatmap of similarity indices of 6 additional markers used to identify major immune lineages. **C**. Heatmap of similarity indices of 4 additional markers to enable deeper phenotyping of B cell subsets, along with the viability dye, and the CD45 antibody-fluorochrome conjugate, giving rise to 30 total markers. Overall panel Complexity Indices, as determined by the Cytek® Full Spectrum Viewer are also included.

**Supplementary Figure 2. Identification of the optimal concentrations for the selected antibody-fluorochrome conjugates.**

Human PBMCs were stained with a given antibody-fluorochrome conjugate at 6-7 different concentrations (1:50, 1:100, 1:200, 1:400, 1:800, 1:1600; for APC-CD25 and Alexa Fluor 700-CD4, 1:25* was also included). Pseudocolor contour plots of live cellular events (pre-gated on live, single cells identified via FSC-A vs. FSC-H) are depicted for each antibody-fluorochrome conjugate. Each contour plot depicts the detector corresponding to the peak fluorochrome emission (Y-axis) and the staining pattern for each concentration (X-axis; concentration decreases from left-to-right). The “optimal” concentration is depicted using a vertical rectangle (rectangles encompassing two concentrations indicate that the optimal chosen concentration lies in between the depicted concentrations). Fluorescence intensities exceeding 300,000 are highlighted in red, as intensities beyond this threshold were generally associated with more significant spillover-spreading. Conjugate concentrations resulting in staining intensities of this magnitude were avoided due to increased spillover spreading, provided that there was no substantial impact on the stain index or the proportion of positive events captured.

**Supplementary Figure 3. Comparison of Alternate and Final Reagents Using Single- and Multicolor Samples.**

**A.** Stain index and the percentage of positive events for alternate antibody-fluorochrome conjugates tested on the same donor sample. Color scales indicate the minimum and maximum range for each parameter across all conjugates tested for a given antigen. **B.** Concentration-dependent spillover spreading (introduced from the fluorochrome on the X axis into the fluorochrome on the Y axis) for four fluorochromes (X axis) with high spectral overlap with other fluorochromes included in this panel. The first of the two plots for a given antibody-fluorochrome conjugate is a dot plot overlay of cells stained with increasing concentrations of the conjugate specified on the X-axis. The second of the two plots for a given antibody-fluorochrome conjugate depicts the spillover spreading introduced from this conjugate (X-axis) into the recipient fluorochrome (Y-axis). **C**. Comparison of CCR4, CD56, and SLAMF7 staining using prior panel iterations and the final 30-color panel. Red X’s and green checkmarks indicate tested reagents that were excluded or retained, respectively. Concentrations highlighted in yellow indicate the reagent concentration selected for inclusion in the final panel.

**Supplementary Figure 4. Assessment of spectral overlap and spillover spreading of the finalized panel.**

**A.** Spillover Spreading Matrix (SSM) for the fully stained panel. The spillover spreading value refers to a measure of the spillover spreading introduced by the conjugate in a given row into the conjugate in a given column. **B.** Bivariate plots of select conjugate pairs exhibiting high spillover spreading values continue to demonstrate adequate marker resolution (parent population consists of “singlets” identified via FSC-A vs. FSC-H and without exclusion of “Dead” or CD45^-^ events).

**Supplementary Figure 5. Validation of the sensitivity and specificity of our negative selection approach for CD4^+^ and CD8^+^ T cells in the absence of anti-CD3.**

**A.** Analysis of each of the four multicolor samples (MC 303458, MC 303444, MC 3015036, MC 3014559) from a public high-dimensional spectral flow dataset (OMIP-069) using a conventional T cell gating strategy leveraging CD3.^69^ **B.** Tables illustrating a comparison of the total cell counts obtained by applying different gating strategies to the four multicolor samples from OMIP-069. “Gating as per Panel A” refers to the gating strategy depicted in Panel A (of this figure) up to the depicted CD4 vs. CD8 bivariate plot. “Gating as per Figure 1B” refers to the gating strategy depicted in Figure 1B of the main text, up to the CD4 vs. CD8 bivariate plot. **C**. UMAP projection of live, CD45^+^ events from all four multicolor samples from OMIP-069 colored by the specified markers. Data from each donor were downsampled to 200,000 live, CD45^+^ events per donor (800,000 total live, CD45^+^ events). Scatterplot overlays depict the events captured using the gating strategy depicted in Panel S5A compared to those captured using the gating strategy depicted in Figure 1B.

**Supplementary Figure S6. Longitudinal application of a 30-color spectral flow cytometry panel enables robust identification of diverse peripheral blood immune subsets with minimal batch effects.**

**A.** UMAP contour density projection of live, CD45^+^ cells from the 18 Bridge samples (1.8 million cells total; 100,000 events/donor; 2 Bridge donor samples used per experimental batch) included in these experiments. Batch-normalized expression profiles of the 28 remaining panel markers is depicted, overlaid on the UMAP embedding. **B.** Concatenated UMAP density scatterplot and corresponding UMAP coordinate density histograms for the specified Bridge sample(s), spanning all nine experimental batches. **C.** Mean Earth Mover’s Distances (EMD)^70^ between UMAP coordinate distributions, calculated either across experimental batches within a given Bridge sample or across Bridge samples within a given experimental batch. Data are depicted as mean ± S.D. *, **, ***, **** = *p* < 0.05, 0.01, 0.001, or 0.0001, respectively, determined via Brown-Forsythe and Welch ANOVA with Dunnett’s T3 multiple comparison correction.

**Supplementary Figure S7. Comparable panel performance on the Aurora CS enables simultaneous high-dimensional phenotyping and purification of peripheral blood immune cell subsets.**

All cytometric data collected on the Cytek Aurora Analyzer was also collected on the Cytek Aurora CS. **A.** Unsupervised, row-clustered heatmap depicting the column z-score for each immune subset quantified in the main text for all 184 donor samples (92 samples per instrument). **B**. Select sorted populations of select samples were reanalyzed on the Cytek Aurora CS to define the post-sort purity.

**Supplementary Figure S8. Associations between clinical and serologic variables among patients with SSc.** Spearman correlation matrix of clinical and serologic variables among patients with SSc (n=37). Statistically significant correlations are indicated with a solid black frame and at least one asterisk. *, **, *** = unadjusted *p* < 0.05, 0.01, or 0.001, respectively, based on Spearman’s rank correlation coefficient.

**Supplementary Figure S9. Additional analyses of innate immune cell subsets.**

**A.** Quantified proportions of CX3CR1^+^ monocyte subsets stratified by disease status and/or sex, shown as boxplots. *, **, ***, or **** = adjusted *p* < 0.05, 0.01, 0.001, or 0.0001, respectively, via Kruskal-Wallis test with Benjamini-Hochberg adjustment for multiple comparisons. **B.** Spearman correlation matrix of clinical features and broad peripheral immune cell subsets. Statistically significant correlations are indicated with a solid black frame and at least one asterisk. *, ** = unadjusted *p* < 0.05 or 0.01, respectively, based on Spearman’s rank correlation coefficient.

**Supplementary Figure S10. Correlation structure of CD8^+^ T cell subsets among patients with SSc.** Spearman correlation matrix of CD8^+^ T cell subsets among patients with SSc.

Statistically significant correlations are indicated with a solid black frame and at least one asterisk. *, **, *** = unadjusted *p* < 0.05, 0.01, or 0.001, respectively, based on Spearman’s rank correlation coefficient.

**Supplementary Figure S11. CD4^+^ T cell subset relationships within the CD4^+^ compartment and across CD8^+^ T cell subsets. A.** Quantified proportions of CD4^+^ T cell subsets not included in the main text, stratified by disease status and/or sex, shown as boxplots. *, **, ***, or **** = adjusted *p* < 0.05, 0.01, 0.001, or 0.0001, respectively, via Kruskal-Wallis test with Benjamini-Hochberg adjustment for multiple comparisons. **B.** Spearman correlation matrix of CD4^+^ T cell subsets. **C.** Spearman correlation of CD4^+^ and CD8^+^ T cell subsets. Statistically significant correlations are indicated with a solid black frame and at least one asterisk. *, **, *** = unadjusted *p* < 0.05, 0.01, or 0.001, respectively, based on Spearman’s rank correlation coefficient. **D.** Scatterplots depicting relationships between selected variable pairs from panel C.

**Supplementary Figure S12. B cell subset relationships within the B cell compartment and across CD4^+^ T cell subsets.**

**A.** Spearman correlation matrix of B cell subsets. **B.** Spearman correlation matrix of B cell and CD4^+^ T cell subsets. Statistically significant correlations are indicated with a solid black frame and at least one asterisk. *, **, *** = *p* < 0.05, 0.01, or 0.001, respectively based on Spearman’s rank correlation coefficient. **C.** Scatterplots depicting relationships between select variable pairs from panel B. Winsorized versions are included as a sensitivity analysis to confirm that the observed associations were robust to extreme values.

## ACKNOWLEDGEMENTS

We thank the staff of the Flow Cytometry Core at the Children’s Hospital of Philadelphia (John Lora, Angela Mckenzie, Kevin Liedel, Chloe Davis, Sophia McKee) for assistance with instrument setup and quality control, and the Human Immunology Core at the Perelman School of Medicine at the University of Pennsylvania (Max Eldabbas, Emileigh Maddox, Tanishk Sinha, and Jiayi Shu) for assistance with bridge sample procurement. We also thank Dr. Jeffrey Carlin from the Benaroya Research Institute for his role in developing the biorepository, and Dr. Divij Mathew from the University of Pennsylvania for guidance regarding the Earth Mover’s Distance calculations used to quantify batch effects using UMAP distributions. Finally, we thank Dr. Maria Jaimes from Cytek Biosciences for helpful discussions and technical insights supporting rigorous validation of the cytometry panel used in this study. This work was supported via grants from the National Institutes of Health (R21-AR081588 to M.C.A), the Penn Colton Center for Autoimmunity and the Institute for Immunology and Immune Health (to M.C.A. and N.J.), the Rheumatology Research Foundation Scientist Development Award (to N.J.), the Scleroderma Research Foundation Postdoctoral Fellowship (to N.J.), and the Penn Rheumatology H. Ralph Schumacher Rheumatology Research Fund (to N.J.).

## AUTHOR CONTRIBUTIONS

Conceptualization: N.J., M.C.A Methodology: N.J.

Data Acquisition: N.J., S.E.P., J.H.B. Data Curation: N.J.

Data Analysis: N.J.

Validation and Visualization: N.J., N.V-P. Consultation/Advice: F.T., J.B.M. Manuscript Draft: N.J.

Manuscript Review and Editing: N.J., F.T., N.V-P., J.B.M., S.E.P., J.H.B., and M.C.A Funding Acquisition: N.J. and M.C.A.

## DATA AVAILABILITY STATEMENT

Raw and/or spectrally unmixed data (.fcs files) will be made available upon reasonable request to the corresponding authors. The catalog number and concentrations of the staining reagents, including the antibody-fluorochrome conjugates used in this panel, are included in the Supplementary Tables.

